# Complete genome sequence and evolution analysis of *Psychrobacter* sp. YP14 from Gammaridea Gastrointestinal Microbiota of Yap Trench

**DOI:** 10.1101/578518

**Authors:** Wei Jiaqiang, Gao Zhiyuan, Liu Whenfeng, Chen Hao

## Abstract

*Psychrobacter* sp. YP14, a moderately psychrophilic bacterium belonging to the class Gammaproteobacteria, was isolated from Gammaridea Gastrointestinal Microbiota of Yap Trench. The strain has one circular chromosome of 2,895,311 bp with a 44.66% GC content, consisting of 2333 protein-coding genes, 53 tRNA genes and 9 rRNA genes. Four plasmids were completely assembled and their sizes were 13,712 bp, 19711 bp, 36270 bp, 8194 bp, respectively. In particular, a putative open reading frame (ORF) for dienelactone hydrolase (DLH) related to degradation of chlorinated aromatic hydrocarbons. To get an better understanding of the evolution of *Psychrobacter* sp. YP14 in this genus, six *Psychrobacter* strains (G, PRwf-1, DAB_AL43B, AntiMn-1,P11G5, P2G3), with publicly available complete genome, were selected and comparative genomics analysis were performed among them. The closest phylogenetic relationship was identified between strains G and K5 based on 16s gene and ANI (average nucleotide identity) values. Analysis of the pan-genome structure found that YP14 has fewer COG clusters associated with transposons and prophage which indicates fewer sequence rearrangements compared with PRwf-1. Besides, stress response-related genes of strain YP14 demonstrates that it has less strategies to cope with extreme environment, which is consistent with its intestinal habitat. The difference of metabolism and strategies coped with stress response of YP14 are more conducive to the study of microbial survival and metabolic mechanisms in deep sea environment.

## 1 INTRODUCTION

The *Psychrobacter* genus, as Gram-negative bacteria most of which are spherical or rod-shaped, heterotrophic and psychrophilic, is ubiquitous in the diverse environments around the world. This genus shows tremendous potential for low-temperature applications, biodegradation especially. (Ayala-del-Rio et al. 2010; Bowman 2006; Bozal et al. 2003). Studies have shown that the content of neutral lipids and phosphate esters in the membrane lipids of the cold bacteria can keep the membrane lipids in a liquid crystal state at low temperatures, which is very important for the survival of cold bacteria at low temperatures (Neidleman,1990; Russell and Fukunaga, 1990; Allen et al,1999). In this genus, many strains have psychrophilic enzymes. Studies on the structure of amylase, subtilisin, and phosphatase have shown that they have better flexibility at low temperatures compared with similar medium-temperature enzymes and high-temperature enzymes (Feller and Gerda, 1997). Therefore, many special psychrophilic enzymes present in this genus have important application value. The habitats of the genus are diverse, and they are isolated from Antarctic soil and seawater, deep-sea, Siberia permafrost. (Bowman et al. 1997; Ayala-del-Rio et al. 2010; Denner et al. 2001; Vishnivetskaya et al. 2000; Xuezheng et al. 2010). However, The size of the genome varies greatly with different species. For example, the genome size of the *Psychrobacter* sp. TB67 is 3.58 MB (Maria Cristiana Papaleo et al. 2012), whereas the genome size of *Psychrobacter* sp. YP14 is only 2.89 MB, which is 80% of the former. Different habitats and genome sizes indicate that the member of this genus has various categories to cope with stress response. Current psychrophilic research focuses on bacteria in the Antarctic and Arctic, and less research on deep-sea psychrophiles. The new member *Psychrobacter* sp. YP14 of *Psychrobacter* genus will attribute to studying on evolution of intestinal microbes in the deep sea.

## 2 MATERIAL AND METHOD

### 2.1 DNA sample preparation

*Psychrobacter* sp. YP14 was isolated in our laboratory from Gammaridea Gastrointestinal Microbiota of Yap trench. The cells of *Psychrobacter* sp. YP14 were cultivated in marine 2216E medium (peptone, 5g; yeast extract, 1g; natural seawater 1L) at 18 °C. The genomic DNA of YP14 was extracted by the SDS method. The prepared DNA was subjected to quality control by agarose gel electrophoresis and quantified by Qubit 2.0 fluorometer (Thermo Fisher Scientific, Waltham, MA, USA).

### 2.2 Genome sequencing

The genome of *Psychrobacter* sp. YP14 was sequenced by Single Molecule, Real-Time (SMRT) technology. Sequencing was performed at Beijing Novogene Bioinformatics Technology Co., Ltd. (Beijing, China). Sequencing data was generated with a PacBio RSII instrument (Pacific Biosciences, Menlo Park, CA, USA). The low-quality reads were filtered by the SMRT Link v5.0.1 and the filtered reads were assembled to generate one contig without gaps.

The coding genes were predicted by GeneMarks (Besemer et al., 2001). Gene islands were predicted with IslandViewer (Bertelli et al., 2017). The tRNAs, rRNAs and sRNAs were predicted by tRNAscan-SE (Lowe and Eddy, 1997), rRNAmmer (Lagesen et al., 2007) BLAST against the Rfam database (Gardner et al., 2009), respectively. A whole genome BLAST search (E-value less than 1e-5, minimal alignment length percentage than 40%) was performed against GO (GeneOntology) (Ashburner et al., 2000), KEGG (KyotoEncyclopedia of Genes and Genomes) (Kanehisa et al., 2006), COG (Clusters of Orthologous Groups) (Tatusov et al., 2003), NR (Non-Redundant Protein Databases databases) (Li et al., 2002), TCDB (Transporter Classification Database) (Saier et al., 2014), Swiss-Prot and TrEMBL (Magrane and Consortium,2011).

### 2.3 Genome evolution analysis

Complete genome sequences of *Psychrobacter* sp. YP14 has been deposited in GeneBank under accession number NZ_CP029789. Another six strains with Complete Genome Assembly level, *Psychrobacter* sp. PRwf-1 (NC_009524), *Psychrobacter* sp. G (NC_021661), *Psychrobacter* sp. P2G3 (NZ_CP012529), *Psychrobacter* sp. P11G5 (NZ_CP012533), *Psychrobacter* sp. AntiMn-1 (NZ_CP012969), *Psychrobacter* sp. DAB_AL43B (NZ_LT799838), were selected and comparative genomics analyses were performed. The evolutionary distances among the seven strains were assessed based on 16S rRNA sequences. The 16sRNA gene sequences were aligned and a maximum likelihood phylogenetic tree was constructed by Mega 7.0 (Kumar et al. 2008). The reliability of the tree was analyzed using bootstrap probabilities. Genome comparisons were performed using the “progressive alignment” option available in the MAUVE program (version 2.4.0). Default scoring and parameters were used for generating the alignment. The circular plot of whole genome identity comparison was generated using BLAST Ring Image Generator (BRIG) (Alikhan et al. 2011) with default parameters. The pan-genome structure was analyzed using Roary (Page et al. 2015) with an identity cut-off of 50%.

## 3 RESULT

### 3.1 General genomic characteristics of *Psychrobacter* sp. YP14

The general genome features of strain *Psychrobacter* YP14 are summarized in Table 1. The sequencing generating 160559 reads with mean read length 7595 bp, totaling 1,219,441,805 bp. The sequencing generated a data volume of 421-fold coverage of the genome and can be assembled into a high quality circular chromosome. The size of the whole genome consisting of a circle chromosome of 2,817,424 bp with 44.66% G + C content. Besides, it owns 4 plasmids. 2,333 coding sequences (CDSs) were predicted, accounting for 83.74% of the whole genome. Among the 2333 CDSs of *Psychrobacter* YP14, 1898 CDSs were classified into COG categories. Among the CDSs with known functions, 53 tRNA and 9 rRNA were detected. Meanwhile, 9 gene islands were identified by the software IslandPath-DIOMB (version 0.2). According to the gene annotation, dienelactone hydrolase (DLH), a key enzyme in the degradation process of chlorinated aromatic hydrocarbon compounds, was predicted in the genome.

**Table 1.**
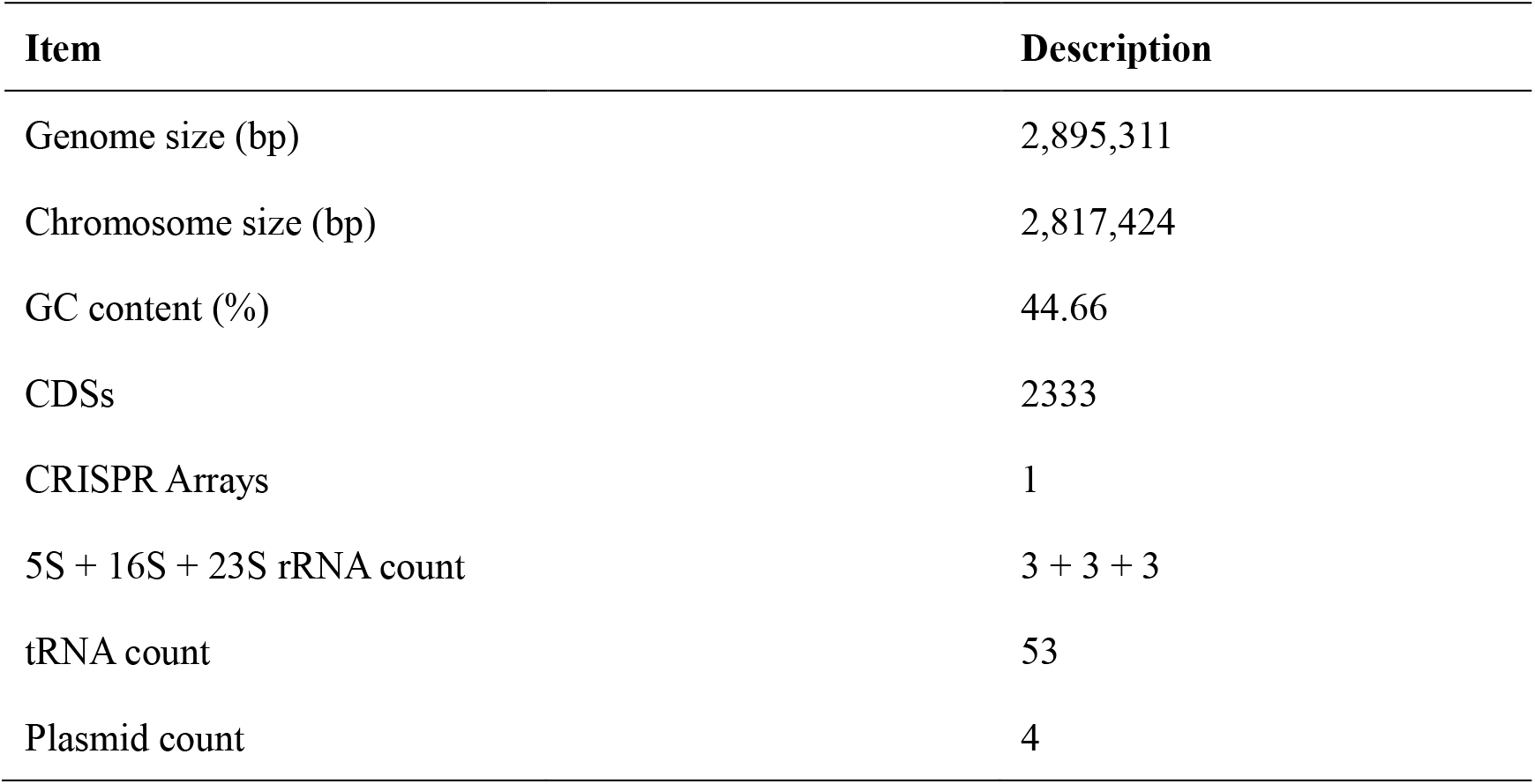
Genomic features of *Psychrobacter* sp. YP14.

### 3.2 Phylogenetic analysis

A summary of the seven strains’ genome features is given in Table 2. Their habits range from the Arctic Ocean, the Pacific Ocean to the Antarctic in the south. Besides, the habit types vary from marine fish, permafrost, deep-sea sediment, tunicates ascidians, to Gammaridea Gastrointestinal. The phylogenetic relationship between the seven strains were analyzed based on 16 S rRNA sequences (Fig 1). The strain YP14 is closest to PRwf-1 in phylogenetic relationship. And the two strains were distant from another five strains.

**Table 2.** Genome features of the seven *Psychrobacter* strains.

| Name | Habitat | Genome<br>Size (bp) | GC<br>(%) | CDS<br>Count | No. of<br>Plasmid | Accession ID |
| --- | --- | --- | --- | --- | --- | --- |
| *Psychrobacter*<br>sp. YP14 | Gammaridea<br>Gastrointestinal Microbiota<br>of Yap trench | 2,895,311 | 44 | 2333 | 4 | NZ_CP029789 |
| *Psychrobacter*<br>sp. PRwf-1 | Host-associated, Skin, Off<br>the coast, northeastern<br>Puerto Rico | 2,995,049 | 45 | 2402 | 2 | NC_009524 |
| *Psychrobacter*<br>sp. G | King George island,<br>Antarctica | 3,113,999 | 42 | 2622 | 3 | NC_021661 |
| *Psychrobacter*<br>sp. P2G3 | Host-associated, Tunicates<br>ascidians, Marine, Arctic | 3366800 | 42 | 2743 | 3 | NZ_CP012529 |
| *Psychrobacter*<br>sp. P11G5 | Host-associated, Tunicates<br>ascidians, Marine, Arctic | 3519382 | 42 | 2859 | 7 | NZ_CP012533 |
| *Psychrobacter*<br>sp. DAB_AL43B | Permafrost in Norway | 3,309,148 | 42 | 2761 | unknown | NZ_LT799838 |
| *Psychrobacter*<br>sp. AntiMn-1 | A deep-sea sediment,<br>Pacific Ocean | 3,138,120 | 44 | 2633 | unknown | NZ_CP012969 |

**Fig.1.**
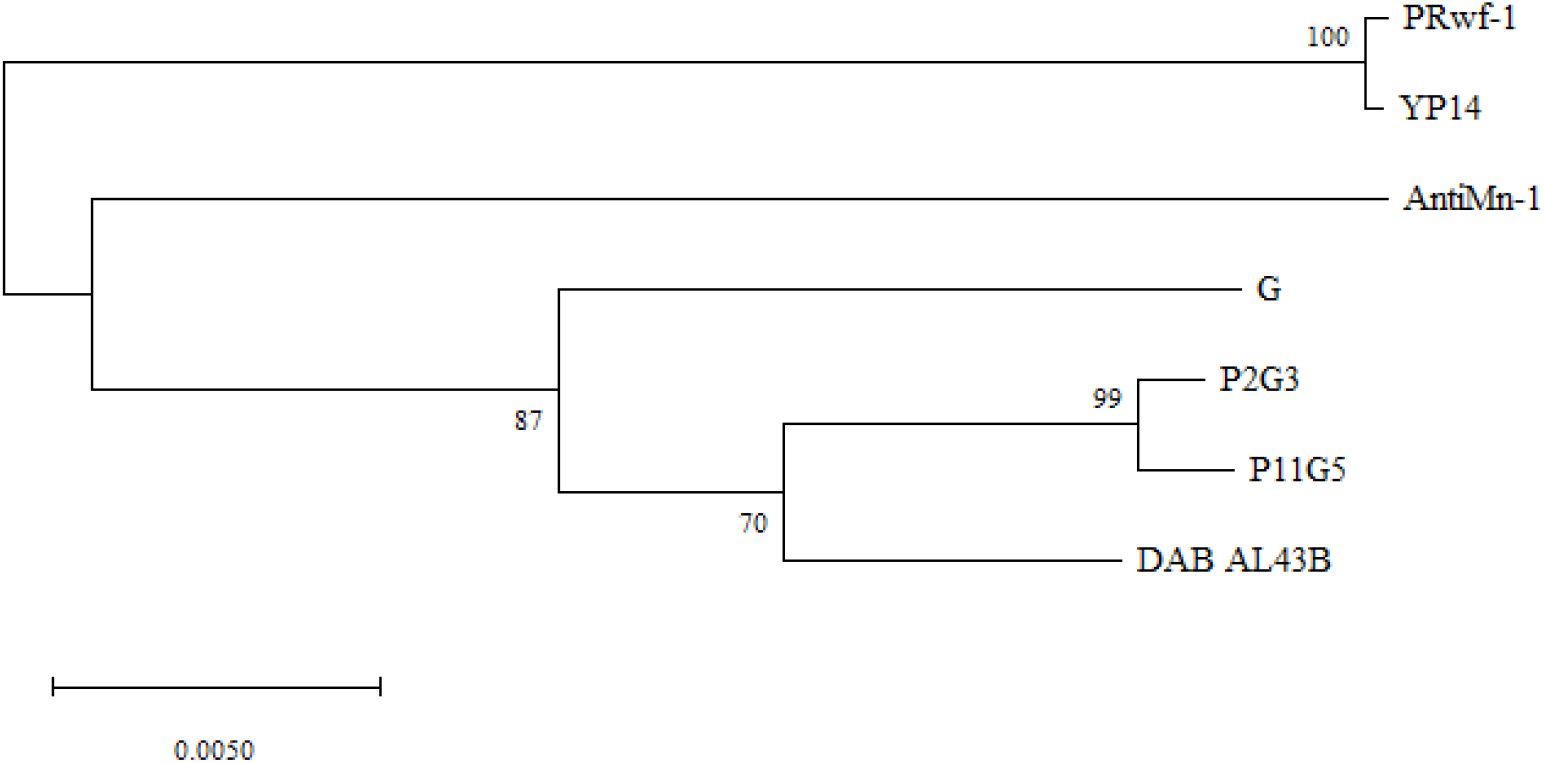
The phylogenetic tree was constructed based on 16S rRNA sequences.

### 3.3 Multi-genome alignment of the seven *Psychrobacter* strains

Complete genome alignments were performed by Mauve and BRIG to show the similarity between seven genomes (Fig 2, 3). The strains YP14 and PRwf-1 show high sequence similarity and numerous locally collinear blocks (LCBs). But several sequence insertion/deletion were observed. Although no inversions were identified, several rearrangements were observed between YP14 and PRwf-1. Besides, P11G5 and P2G3 reveals high sequence similarity.

**Fig.2.**
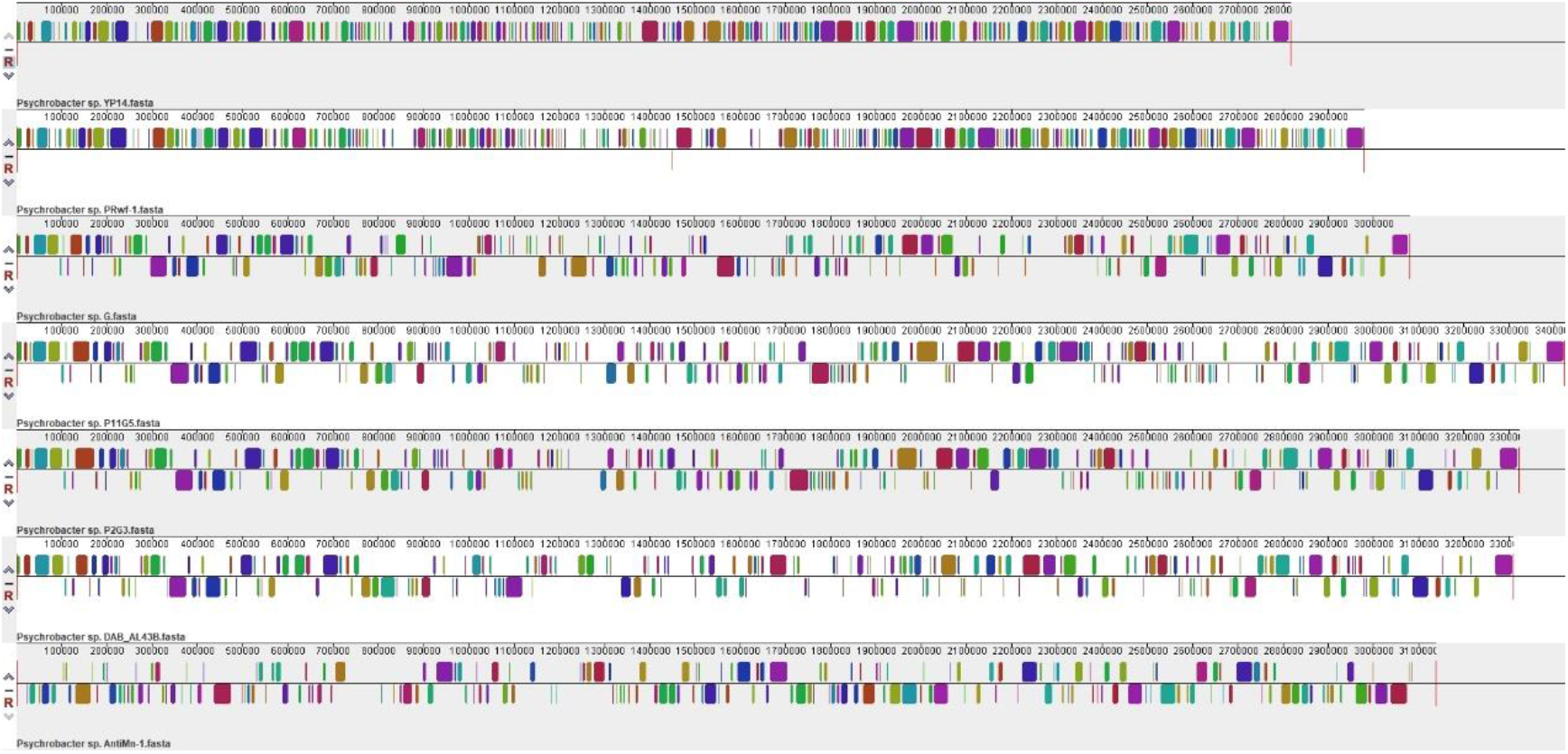
Mauve alignment of seven *Psychrobacter* strains.

**Fig.3.**
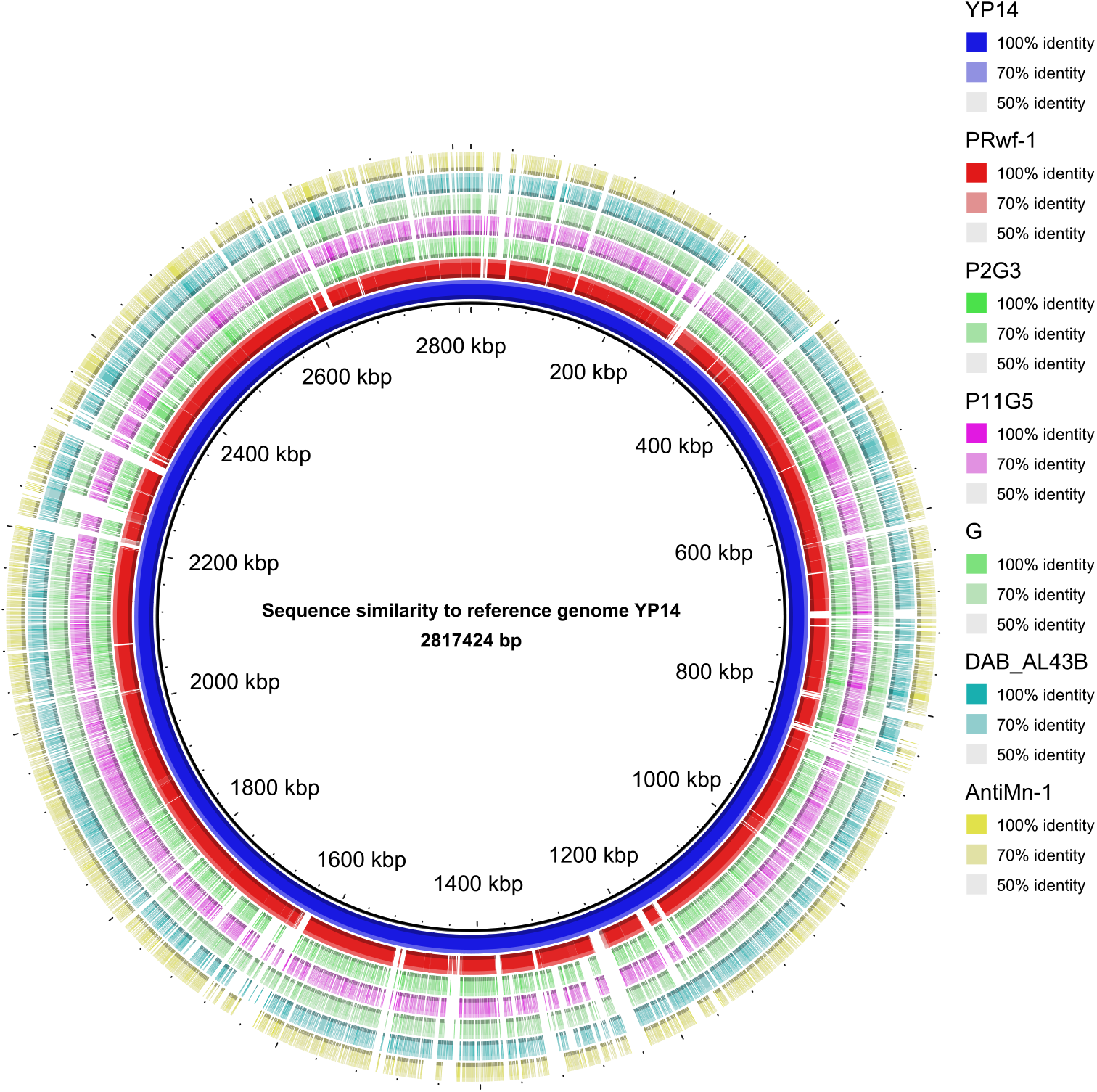
Sequence similarity was calculated with BlastN.

### 3.4 Pan-genome structure of the seven strains

The pan-genome structure of the seven strains was analyzed to get their variations in gene contents (Fig 4). A total of 5307 gene clusters were identified, and 1516 (28.5%) core genes were identified from all of them (Fig 4.). Unique gene varies greatly among different strains. The number of unique gene clusters in strain YP14 is 124, which is the least of the seven strains. The numbers of unique gene clusters of stain PRwf-1, G, P2G3 and P11G5 are only 251, 248, 139 and 157 respectively, however it can reach 504 and 538 in strain AntiMn-1 and DAB_AL43B, respectively. (Fig 4).

**Fig.4.**
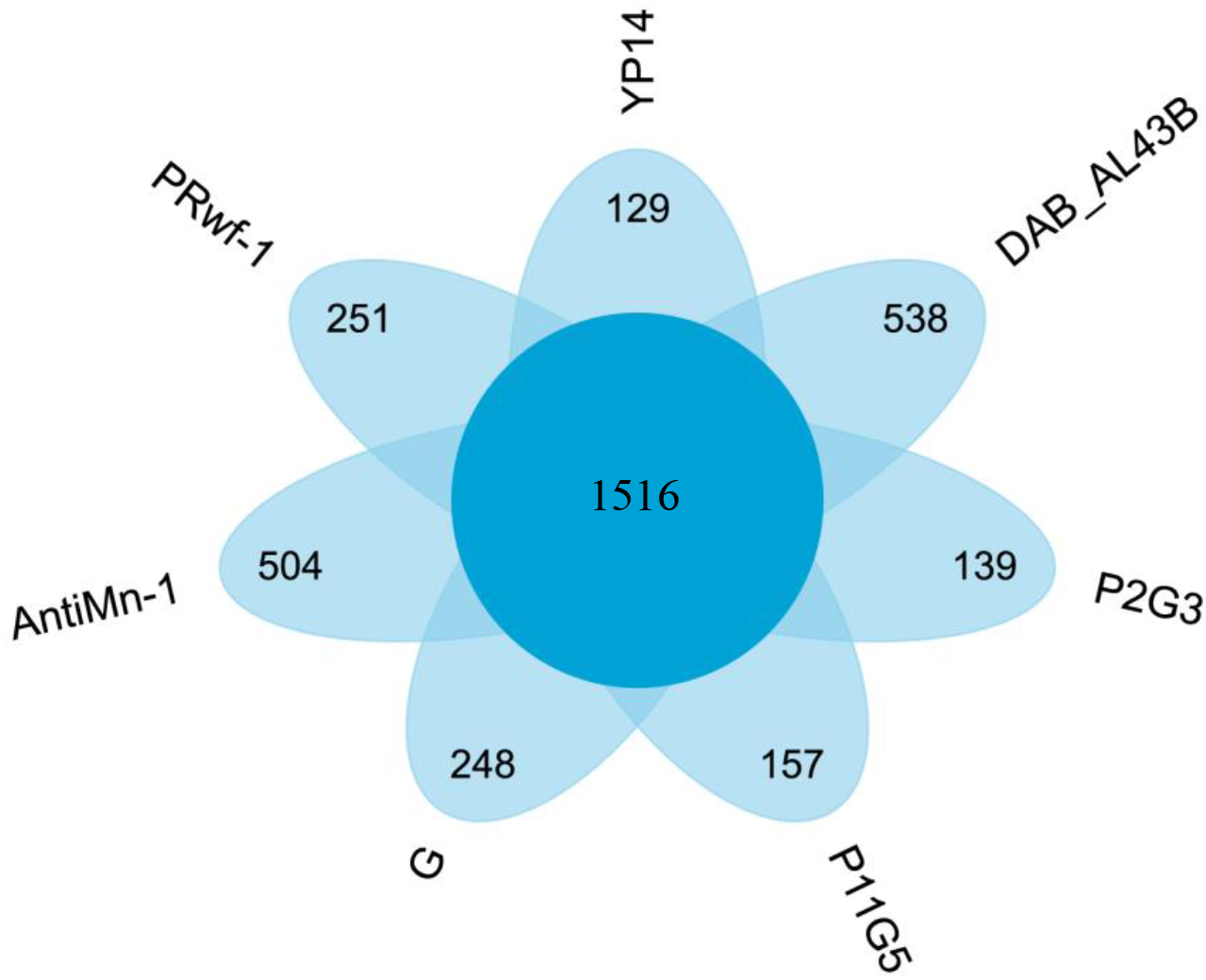
The number of strain-specific genes in seven *Psychrobacter* strains.

The COG assignment of the chromosomes, core genes, and unique genes in the seven strains is given in Fig 5. The seven strains’ chromosome genes showed similar COG distribution profile. The number of genes related to [K] Transcription in strain PRwf-1’s (91) and YP14’s (81) chromosome is lower than the other five strains. Besides, the number of genes related to [U] Intracellular trafficking, secretion, and vesicular transport in strain PRwf-1’s (29) and YP14’s (25) chromosome is lower than the other five strains. Of the 1516 core genes clusters, 207 encode poorly characterized proteins. The most enriched COG categories are [C] Energy production and conversion (126), [E] Amino acid transport and metabolism (153) and [J] Translation, ribosomal structure and biogenesis (194). The number of COG clusters related to [X]Mobilome: prophages, transposons in strain YP14 (33) is lower than in strain PRwf-1 (89).

**Fig. 5.**
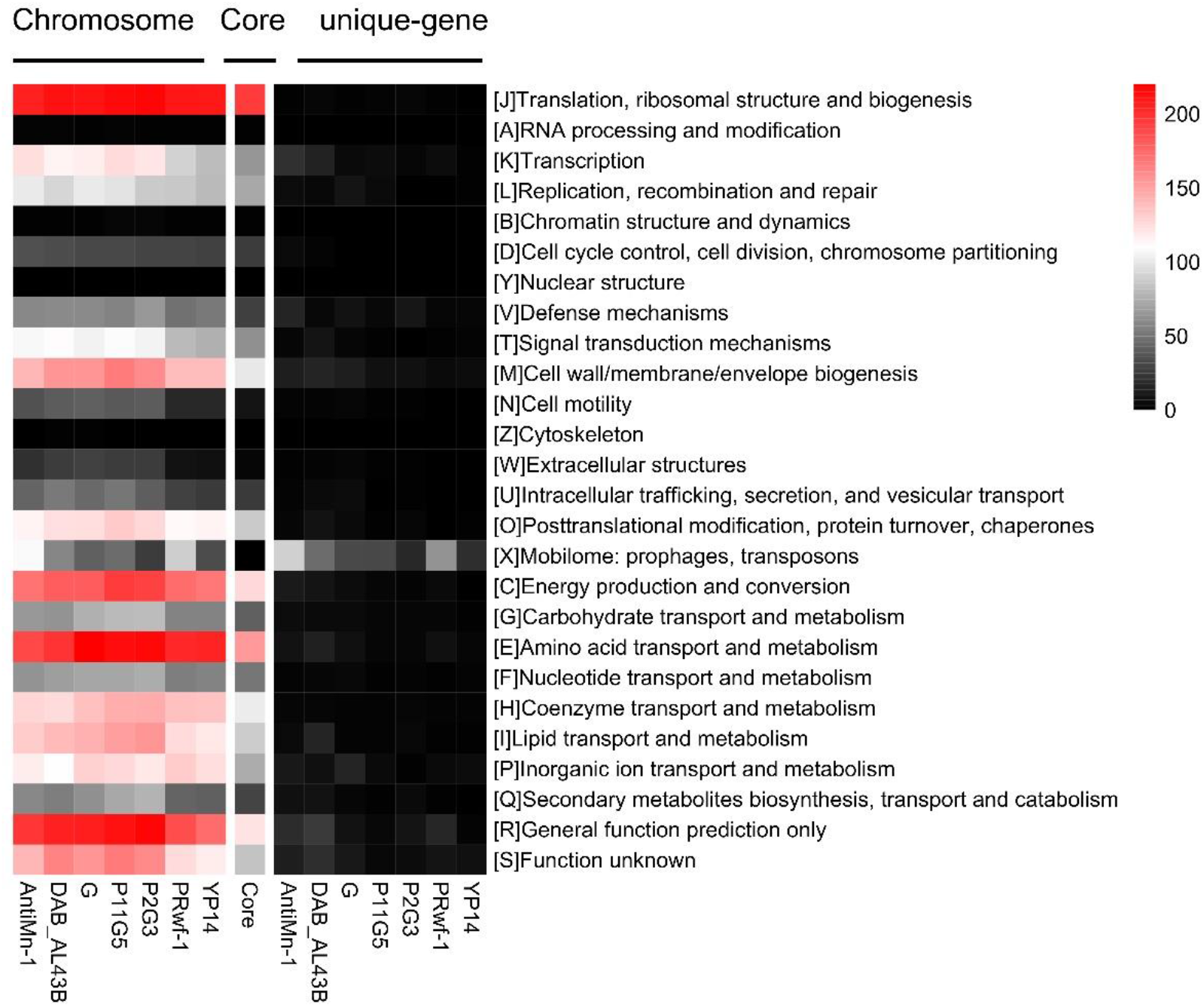
COG assignment of chromosome, core, unique genes for the seven genomes.

Genes associated with stress response were identified from the seven genomes and are given in Fig 6. The stress factors are temperature, osmotic, oxidative, starvation. The choline–glycine betaine transporter is the most abundant osmotic regulator in these seven strains. Trehalose-6-phosphate synthase and trehalose-6-phosphate gene clusters are absent in strain YP14 and PRwf-1’s genome. Besides, the SOS-response transcriptional repressors are absent in strain YP14’s genome. Several protein families associated with oxidative stress response were identified from seven genomes. They were three peroxiredoxin families, two superoxide dismutase and one catalase family, respectively. Three starvation-related protein families were identified from the each of the seven genomes.

**Fig.6.**
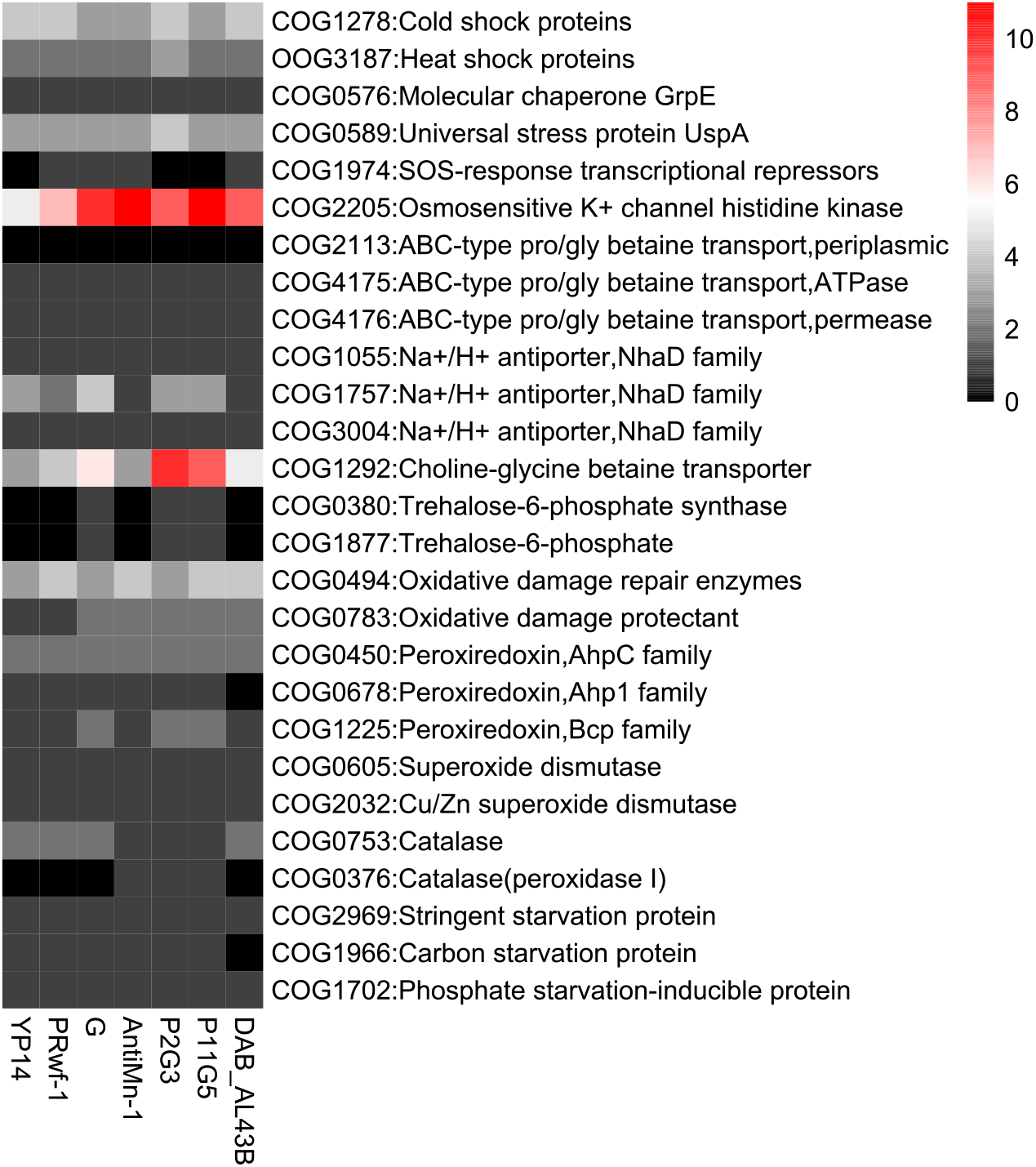
Abundance of stress response-related genes from the seven genomes.

## 4 DISCUSSION

Genome sequencing of the strain YP14 predicted a putative open reading frame (ORF) for dienelactone hydrolase (DLH) with 74.3% sequence similarity to *Psychrobacter* sp. G. Dienelactone Hydrolase is a key enzyme in the degradation of chlorinated aromatic hydrocarbons. Chlorinated aromatic hydrocarbons are common environmental pollutants such as the representative insecticides 2,4-D (2,4-dichlorophenoxyacetate) and 3-CB (3-chlorobenzoate). During the decomposition process, 2,4-D and 3-CB are first decomposed into chlorocatechols, and then degraded successively by four enzymes (chlorocatechol-1,2-dioxygenase, chloromuconate cycloisomerase, DLH and maleylacetate reductas). Among them, DLH is a key enzyme for decomposing diene lactones and their analogues. Due to many proteins isolated from deep-sea bacteria are very different from other habits. Several cold-adapted protein characteristics like glycine clustering in the binding pocket, reduced protein core hydrophobicity, as well as the absence of proline residues in loops are exhibited by the proteins. The identification of DLH in YP14 would attribute to the degradation of pollutants at low temperatures.

Multi-genome alignment of the seven genomes showed high identity and numerous sequence synteny between YP14 and PRwf-1 reveal their close phylogenetic relationship. This is consistent with the results of the phylogenetic tree and ANI values. The different location of YP14 and PRwf-1 were isolated showed that gene exchange is more likely to occur in environments with higher bacterial population density.

Through phylogenetic analysis, we found that YP14 and PRwf-1 have very close evolutionary origins. The Average Nucleotide Identity (ANI) reach 98.56%. Based on Konstantinidis’ statements, two genomes that show higher than 95% ANI are belonging to the same species (Konstantinidis et al. 2006). Therefore, PRwf-1 and YP14 were suggested that they belong to the same species.

COG cluster analysis of core genes, chromosomes, and unique genes showed differences in function between strain YP14 and other six *Psychrobacter* strains. In strain PRwf-1’s unique-gene, there are 64 genes of prophages and transposons (25%) (Fig. 5), much higher than the strain YP14 (20), which means that strain PRwf-1 may undergo frequent gene gain and loss for adaptation to its distinct niche. Because of YP14’s relatively stable living environment in the intestinal tract, its gene exchange mostly occurs between intestinal microorganisms. Thus, compared with other six strains, YP 14 has fewer gene mutations and exchanges.

The identification of stresses related genes revealed that at least three Cold shock proteins (CSPs) in the seven genomes. The universal existence of CSPs may help *Psychrobacter* strains adapt to the frequently encountered cold stresses (Song et al. 2012). Besides, the absence of the SOS-response transcriptional repressors indicated that the intestinal YP14 less exposure to DNA damage. Trehalose is both a storage saccharide and an important product of stress metabolism. It has the function of protecting biological cells and biologically active substances from damage under adverse environmental conditions such as dehydration, drought, high temperature, freezing, high osmotic pressure and toxic reagents. Trehalose-6-phosphate synthase is a key enzyme in the synthesis of trehalose. We found strain YP14 absent Trehalose-6-phosphate synthase, indicating that strain YP14 has less stress pressure in intestine. The distribution of different osmotic regulators in these 7 genomes indicates that these strains have adopted a variety of strategies to fit with changes in osmotic pressure.

## 5 CONCLUSION

The unique habitat of the deep sea makes microbes have different functions from bacteria in other places. Despite different geographical locations and hosts, the strain YP14 and PRwf-1 showed the closest phylogenetic relationship based on 16s gene and ANI values. Whole genome alignments with MAUVE and BRIG revealed the presence of numerous blocks of homologous regions. The results indicated that the two strain (YP14 and PRwf-1) with distant geographical distance (Puerto Rico and Yap Trench) showed the closest phylogenetic relationship and highest sequence similarity. The pan-genome structure of seven strains showed the fewest unique gene in strain YP14. The COG cluster gene associated with transcription, Intracellular trafficking, secretion, and vesicular transport on the chromosomes of the two strains is less than the other five strains, which indicated that they may have weaker metabolic abilities. High differences in genes related to stress response suggest that stress response mechanisms employed by members of this genus may be highly diversified. Compared with PRwf-1, which was isolated from fish skin, the strain YP14 with lower sequence rearrangements were isolated from Gammaridea Gastrointestinal. YP14 has fewer COG clusters associated with transposons and prophage. The absence of related trehalose synthesis and SOS-response transcriptional repressors, showing that YP14 has fewer strategies for dealing with changes in osmotic pressure and other extreme conditions. In addition, there are fewer Osmosensitive K+ channel histidine kinase genes, which also means that its habit make it unnecessary to face strong osmotic pressure changes. Compared with other *Psychrobacter* strains, YP14 has fewer strategies fit with stress response and this is the result of it’s adaptation to the environment.

## 6 DATA AVAILABILITY STATEMENT

All sequence data is available at NCBI (https://www.ncbi.nlm.nih.gov/genome/). The datasets generated and/or analyzed during the current study are available from the corresponding author on reasonable request.

